# Loss of neurofibromin alters adult metabolism via effects during a developmental critical period

**DOI:** 10.64898/2026.07.29.741559

**Authors:** Catherine Steele, Ryan Weaver, Seth M. Tomchik

## Abstract

Metabolic alterations commonly accompany neurodevelopmental disorders and may contribute to their pathophysiology. Neurofibromatosis type 1 (OMIM 162200) is a genetic disorder that results from mutations in the NF1 gene and its encoded neurofibromin protein (Nf1). The disorder is multisystemic, affecting multiple aspects of development, physiology, and brain function. In addition, recent evidence suggests that Nf1 deficiency alters metabolic function in both humans and animal models. Whether the metabolic alterations result from changes in neurodevelopment is not known. Here we approach this question in *Drosophila melanogaster*, which expresses a conserved *NF1* gene, exhibits phenotypes reminiscent of the human disease, and shares key developmental mechanisms with humans. Flies with *nf1* mutations or RNAi-mediated knockdown exhibit altered metabolism in adulthood. Conditional Nf1 inactivation revealed that the adult metabolic phenotype resulted from loss of Nf1 in neurons during a developmental critical period (third instar larva/pupa), which corresponded to the period of nervous system maturation. Prior to the developmental critical period, *nf1* mutants did not exhibit metabolic difference - rather, the metabolic alterations appeared only after the critical period. This suggests that the adult phenotype results from the onset of the developmental alteration during the critical period. High-resolution respirometry on adult mitochondria revealed no differences in complex I/II function or fatty acid oxidation between *nf1* mutants and controls, suggesting that the metabolic alterations localize upstream of the electron transport chain at the cellular level. Overall, these data suggest that loss of Nf1 alters adult metabolism via effects during a critical period of nervous system development.

## Introduction

Neurofibromatosis type 1 (NF1) is an autosomal-dominant monogenetic disorder that is caused by mutations in the *NF1* gene, which encodes the tumor suppressor protein neurofibromin (Nf1). The Nf1 protein acts as an inhibitory regulator of Ras via a central Ras-GTPase activating protein (GAP)-related domain (GRD). Loss of Nf1 increases activity of Ras and multiple pathways downstream that regulate multiple cellular processes [1–5]. Manifestations of NF1 include nervous system tumors, cognitive and behavioral symptoms, altered brain morphology, bone defects, and café au lait spots [1, 2, 6, 7]. In addition, patients with NF1 show altered metabolism, with symptoms including short stature, lower fasting blood glucose levels, a lower prevalence of diabetes mellitus and increased resting energy expenditure among females with NF1 [8–10]. These features suggest that individuals with NF1 experience shifts in cellular and systemic metabolic function.

Nf1 deficiency alters metabolism in animal models as well as humans. Heterozygous Nf1 mice exhibit increased insulin sensitivity and glucose utilization, as well as reduced fat mass and increased lean mass [11]. In *Drosophila*, mutations in Nf1 result in decreased body size and triglyceride stores, shortened lifespan, increased metabolic rate, and increased rate of lipolysis [12–14]. Neuronal mechanisms play a role in regulating metabolic homeostasis through Nf1 - loss of Nf1 in *Drosophila* exhibit increased metabolic rate and lipid turnover rate, as well as decreased respiratory quotient via neuronal mechanisms [12].

In addition to metabolic changes, neurodevelopmental alterations accompany NF1. The disorder affects multiple aspects of neurodevelopment, including altering nervous system structural features [1, 7, 15]. It is unclear whether/how such anatomical alterations drive the symptoms of the disease. This is a question of critical consequence, as therapeutic strategies to date have focused on drug treatment after nervous system maturation; these have produced limited efficacy thus far on cognitive symptoms [16–20]. A previous study in *Drosophila* found that loss of Nf1 in neurons during a developmental critical period was responsible for Nf1-dependent alterations on a behavioral phenotype (increased grooming) [21]. This suggests that neurodevelopmental expression of Nf1 may be critical for normal adult behavior. Relatedly, cellular signaling pathways downstream of Nf1 exert neurodevelopmental effects in mammals [22]. This raises the question of how loss of Nf1 affects neurodevelopment, and how that in turn affects other symptoms of the disease, such as the metabolic features [1]. Here we approached this question using a combination of in vivo genetic analysis and high-resolution respirometry at the organismal and mitochondrial levels.

## Results

### Increased metabolic rate in *nf1* mutants is due to neuronal effects of Nf1, potentially during neurodevelopment

Nf1 is a major Ras GAP that increases metabolic rate in adult *Drosophila* (Fig. 1A) [12]. To investigate the potential developmental contributions of Nf1 to metabolism, we first quantified metabolic rate with multiple genomic *nf1* mutations (Fig. 1B). CO_2_ production was measured via respirometry in 5-day-old adult flies (Fig 1C). We first tested a large genomic deletion, *nf1*^*P1*^ [14]. *nf1*^*P1*^ mutants were tested against two controls: the parental (w+) K33 line containing the unexcised P element that was used to generate *nf1*^*P1*^ via imprecise excision [23], as well as the wCS10 (w-) line that both *nf1*^*P1*^ and K33 were backcrossed into [24]. CO_2_ production was significantly higher in *nf1*^*P1*^ than both K33 and wCS10 (Fig. 1D). This suggested that Nf1 deficiency increased CO_2_ production and verified that the w+/w-status of the fly line did not affect the results of the assay. Loss of Nf1 increases feeding [12] and significantly enlarges the crop [25], which could affect gut microbiota quantity and/or content. To examine whether this underlies the metabolic effect, we tested the effect of including an antibiotic (streptomycin) in the flies’ food prior to testing metabolism. *nf1*^*P1*^ mutants continued to exhibit significantly increased CO_2_ production relative to controls following streptomycin feeding (Fig. 1E), suggesting that the gut microbiota and bacteria in the food did not significantly impact the Nf1 metabolic effect. The *nf1*^*P1*^ mutation is a large deletion that encompasses most of the Nf1 coding region plus at least two adjacent genes in the Enhancer-of-split complex (E(spl)-C) [14]. Therefore, to further challenge the results from this mutation, we tested two additional mutants. *nf1*^*E1*^ is a nonsense point mutation upstream of the catalytic GRD that truncates the protein after 1061 amino acids [26]. The *nf1*^*E1*^ mutant exhibited higher CO_2_ production relative to its genetic background control (iso2,3) (Fig. 1F). Finally, we tested *nf1*^*C1*^, a small indel in exon 2 (Nf1del162-163), that eliminates protein expression [27]. Like both other genomic mutants, *nf1*^*C1*^ exhibited increased CO_2_ production relative to its genetic background control (iso2,3) (Fig. 1G). Thus, three different *nf1* genomic mutant lines exhibited altered metabolism, as assayed by CO_2_ respirometry.

**Figure 1.**
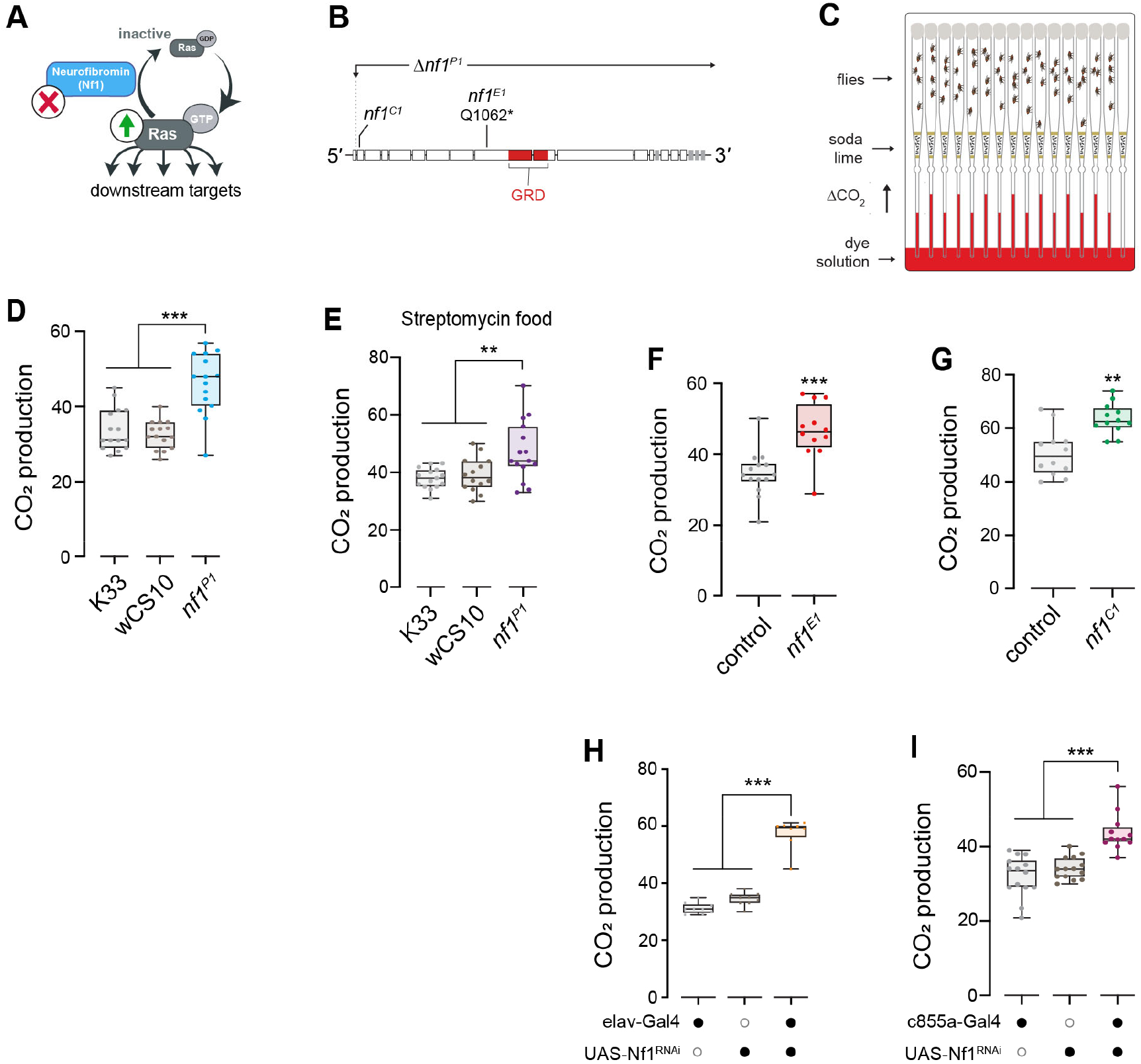
Increased metabolic rate in adult *nf1* mutants is due to neuronal effects of Nf1, potentially during neurodevelopment. **(A)** Diagram of Nf1’s Ras GAP signaling functionality. **(B)** Diagram of the *Drosophila Nf1* gene and the mutations used in this study. GRD: GAP-related domain. **(C)** Diagram of CO_2_ respirometry setup. **(D)** CO_2_ production in *nf1*^*P1*^ mutants, compared to K33 and wCS10 controls. ***p < 0.001 (ANOVA/Dunnett). **(E)** CO_2_ production in *nf1*^*P1*^ mutants fed with streptomycin food, compared to K33 and wCS10 controls. **p<0.01 (ANOVA/Dunnett). **(F)** CO_2_ production in *nf1*^*E1*^ mutants, compared to iso2,3 controls. ***p < 0.001 (Mann-Whitney). **(G)** CO_2_ production in *nf1*^*C1*^ mutants, compared to iso2,3 controls. **p < 0.01 (Mann-Whitney). **(H)** CO_2_ production in flies with Nf1 knocked down pan-neuronally using elav-Gal4 to drive UAS-Nf1 RNAi. ***p < 0.001 (ANOVA/Dunnett). **(I)** CO_2_ production in adult flies with Nf1 knocked down in neuroepithelial cells using c855a-Gal4 to drive UAS-Nf1 RNAi. ***p < 0.001 (ANOVA/Dunnett).

Loss of Nf1 alters metabolism via effects in neurons [12], though it is unknown whether this is due to developmental or adult expression of Nf1. To begin addressing this question, we first compared the CO_2_ production when knocking down Nf1 with RNAi [24, 28] using a pan-neuronal driver and a neuroepithelial driver that expresses during neurodevelopment. Panneuronal knockdown of Nf1 using elav-Gal4 increased CO_2_ production relative to heterozygous Gal4 driver and UAS effector controls (Fig 1H). Next, we knocked down Nf1 in neuroepithelial cells using the c855a-Gal4 driver. This Gal4 line targets the larval neuroepithelium and was selected due to the similarity between the transition of the neuroepithelial to neuroblast in *Drosophila* and the development of the cerebral cortex in mammals [29–31]. When knocking Nf1 down in neuroepithelial cells, a significant increase in CO_2_ production was observed in the experimental group compared to the heterozygous Gal4 and UAS controls (Fig. 1I). This suggested that the increased CO_2_ production in *nf1* mutants, while observed in adults, could involve underlying neurodevelopmental mechanisms.

### Loss of Nf1 alters metabolism via effects during a developmental critical period

To test whether the Nf1-dependent increase in CO_2_ production was due to neurodevelopmental effects, we conditionally knocked down Nf1 pan-neuronally at different developmental time points. Nf1 expression was knocked down with RNAi (UAS-Nf1 RNAi) under control of a pan-neuronal Gal4 driver (elav-Gal4) and a temperature-sensitive Gal4 repressor (tubGal80^ts^) [32] (Fig. 2A). Flies of the experimental genotype (expressing the RNAi) were compared to genetic controls (expressing GFP) that were subjected to the same temperature shifts. Four conditional knockdown protocols were tested: no induction (negative control), early development Nf1 knock-down (embryo through 2^nd^ instar larval stage), late development Nf1 knockdown (3^rd^ instar through pupal stage), and adult-only knockdown (following eclosion) (Fig. 2B). Males and females were tested separately. In the negative control animals (with no induction), there was a difference in baseline CO_2_ production between the genotypes harboring the Nf1 RNAi and those containing GFP (lower in the RNAi animals) (Fig. 2C,D). Thus, this represents the baseline for examining the effects of induction.

**Figure 2.**
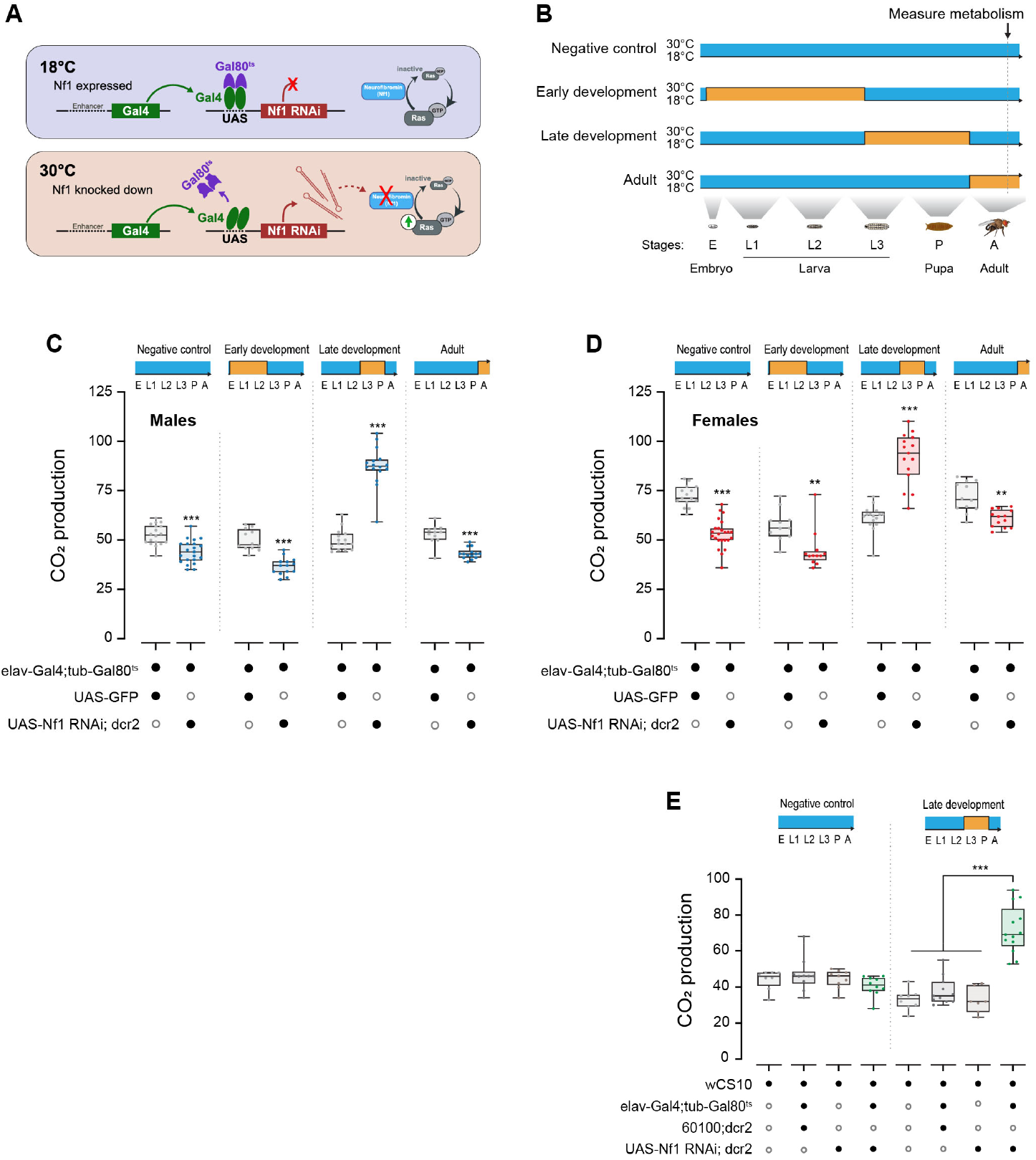
Loss of Nf1 alters metabolism via effects during a developmental critical period. **(A)** Diagram of the Gal80^ts^ conditional knockdown strategy, following [32]. **(B)** Induction protocols to knock Nf1 down during different developmental time windows. **(C)** CO_2_ production in male flies, comparing flies with Nf1 knockdown to GFP-expressing genetic controls. ***p<0.001 (ANOVA/Šidák). **(D)** CO_2_ production in female flies, comparing RNAi-expressing experimental flies to GFP-expressing controls. **p < 0.01, ***p < 0.001 (ANOVA/Šidák). **(E)** CO_2_ production during late development knockdown, compared experimental flies and genetic controls at the permissive (18°C) (negative control) and restrictive (30°C) temperatures. ***p < 0.001 (ANOVA/Šidák).

Knockdown of Nf1 during early development did not alter CO_2_ production in either males (Fig. 2C) or females (Fig. 2D). While there was significantly less CO_2_ production in the Nf1 RNAi group than the GFP-expressing genotype in both cases, it was similar to the uninduced GFP controls and cannot account for the increase in CO_2_ production observed in the *nf1* mutant adults (Fig. 1). In contrast, knockdown of Nf1 in late development produced a significant increase in CO_2_ production in both males (Fig. 2C) and females (Fig. 2D). Adult-selective knock-down of Nf1 did not increase CO_2_ production. As with the early development knockdown, there was significantly less CO_2_ produced by the Nf1 RNAi line compared to the GFP-expressing control, mirroring the uninduced control in both males and females (Fig. 2C,D). To confirm that the increased CO_2_ production observed with Nf1 knockdown in late development was due to the RNAi, we carried out additional genetic controls at that time point. The genetic background line, wCS10, as well as flies heterozygous for each of the genetic elements used in the conditional knockdown experiments, were tested for CO_2_ production in the absence or presence of a late development temperature shift (Fig. 2E). Late development-specific knock down produced a significant increase in CO_2_ production relative to the genetic controls, and there was no effect in uninduced (no temperature shift) flies (Fig 2E). Taken together, these data suggest that Nf1 is required during a developmental critical period to regulate adult metabolic rate in *Drosophila*.

### Metabolic changes first appear in adulthood

Since Nf1 regulated adult metabolism via effects during a developmental critical period, we wondered when the metabolic phenotype emerges during the lifecycle. As a first test, metabolic rate was measured via CO_2_ respirometry in *nf1* mutants (as above, examining *nf1*^*P1*^, *nf1*^*E1*^, and *nf1*^*C1*^). Since the critical period is in late development, we initially focused on the pupal and young adult life stages to determine whether the phenotype had become established at that time point. In late development, at the 48 hr and 96 hr pupal stages, *nf1*^*P1*^ mutants did not exhibit a difference in CO_2_ production relative to wCS10 controls (Fig. 3A). Similarly, there was no increase in CO_2_ production between *nf1*^*E1*^ mutants (Fig. 3B) or *nf1*^*C1*^ mutants (Fig. 3C) relative to their respective controls. Moving into adult animals, there was a detectable, significant increase in CO_2_ production 1 day after pupal eclosion in *nf1*^*P1*^ mutants (Fig. 3D), which was also observed 3 days after eclosion (Fig. 3D) (akin to the 5 day-old flies tested previously [Fig. 1D]). *nf1*^*E1*^ and *nf1*^*C1*^ mutants exhibited qualitatively similar increases in CO_2_ production after eclosion, though the timeline and dynamics differed. *nf1*^*E1*^ mutants did not yet exhibit higher CO_2_ production than controls at 1 day-old, but a difference emerged at 3- and 5-days old (Fig. 3E). In *nf1*^*C1*^ mutants, there was a significant decrease in CO_2_ production relative to controls in 1 day-old adults, no difference at 3 days old, and a significant increase appeared at 5 days after eclosion (Fig. 3F). Integrating all of these findings, CO_2_ production increased beginning in adulthood and was significantly elevated by 5 days post-eclosion in all mutants.

**Figure 3.**
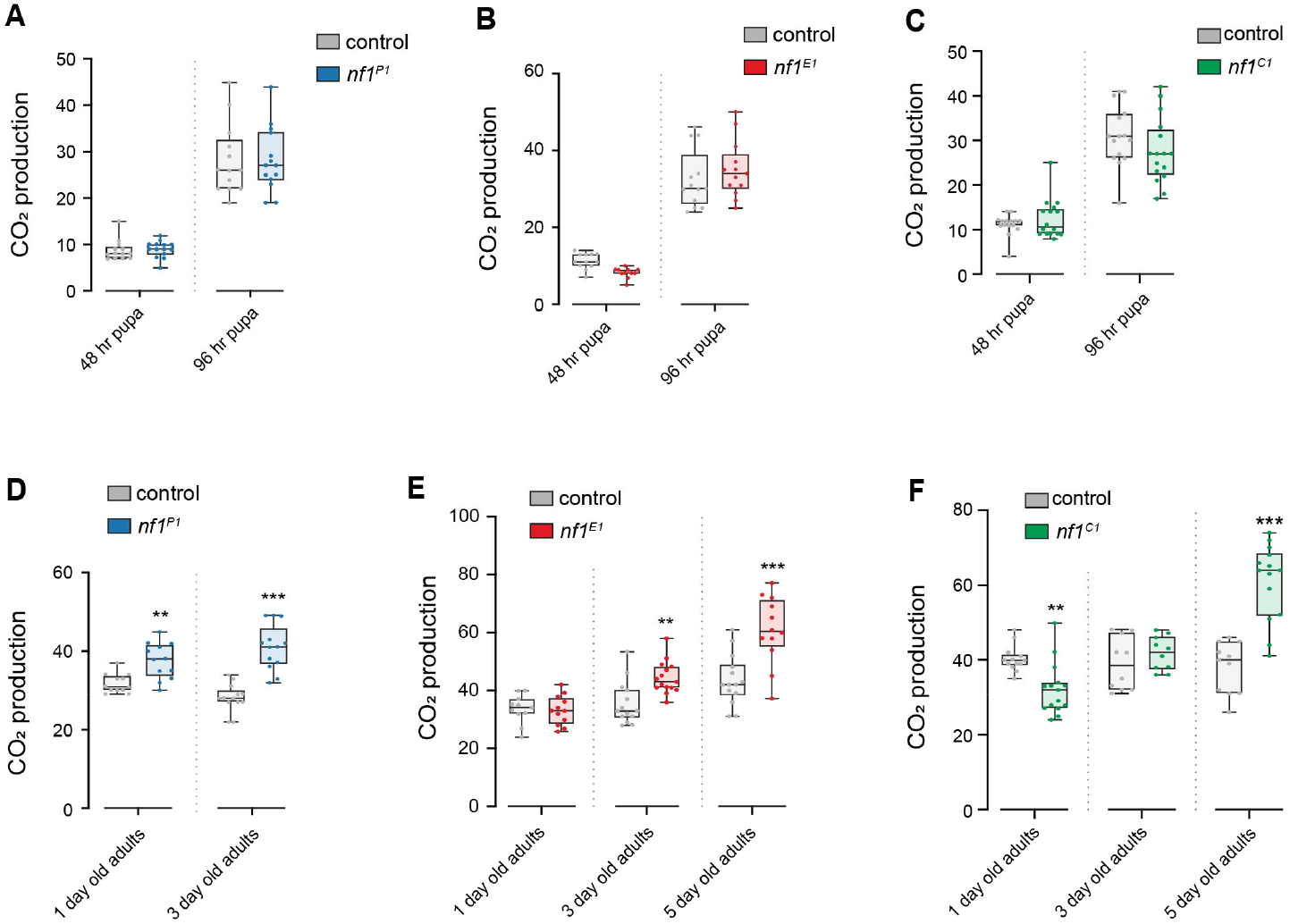
Increased CO_2_ production with Nf1 deficiency appears in adulthood. **(A)** CO_2_ production in *nf1*^*P1*^ mutants during the pupal stage, compared to the control line (wCS10). p = 0.99 (ANOVA/Šidák). **(B)** CO_2_ production in pupal *nf1*^*E1*^ mutants, compared to iso2,3 controls. p = 0.348 for 48 hr, p = 0.461 for 96 hr (ANOVA/Šidák). **(C)** CO_2_ production in pupal *nf1*^*C1*^ mutants, compared to the control line (iso2,3). p = 0.78 for 48hr, p = 0.171 for 96hr (ANOVA/Šidák). **(D)** CO_2_ production in adult *nf1*^*P1*^ mutants (1 day old and 3 day old), compared to wCS10 controls. **p < 0.01, ***p < 0.001 (ANOVA/Šidák). **(E)** CO_2_ production in adult *nf1*^*E1*^ mutants (1 day old, 3 day old, and 5 day old), compared to iso2,3 controls. p = 0.0.998 for 1 day old, **p <0.01, ***p < 0.001 (ANOVA/Šidák). **(F)** CO_2_ production in adult *nf1*^*C1*^ mutants (1 day old, 3 day old, and 5 day old adult flies), compared to iso2,3 controls. **p < 0.01, p = 0.777 for 3 day old, ***p < 0.001 (ANOVA/Šidák).

CO_2_ production and O_2_ consumption are correlated processes, and measurement of each reflects aspects of metabolic rate [12, 33]. To examine the ontogeny of the metabolic phenotype with high resolution, we tested O_2_ consumption rate across development using a microplate respirometry system (Fig. 4A). We first compared *nf1*^*P1*^ mutants to control embryos, observing no significant difference in O_2_ consumption rate (Fig. 4B). Similarly, there was no difference in O_2_ consumption rate between *nf1*^*P1*^ mutants and control 1^st^ instar larvae (Fig. 4C) or 2nd instar (Fig. 4D). In contrast, 3^rd^ instar larvae exhibited an increase in O_2_ consumption relative to controls (Fig. 4E). Moving into the pupal stage, we examined O_2_ consumption in *nf1*^*P1*^ mutants across multiple time points – 0, 24, 48, 72, and 96 hr after pupal formation. There was a significant increase in O_2_ consumption in *nf1*^*P1*^ mutants at 0 hr after pupal formation, but none at any of the other pupal time points (Fig. 4F). Finally, we examined O_2_ consumption in adults. *nf1*^*P1*^ mutants exhibited significant increases in O_2_ consumption rate relative to controls at 1, 3, and 5 days post-eclosion (Fig. 4G). This mirrored the results with CO_2_ consumption in these mutants (Figs. 1,3). Overall, we saw an increase in O_2_ consumption around the time of the Nf1 critical period (3^rd^ instar larvae and early pupal phase), and then an increase in O_2_ consumption again following eclosion.

**Figure 4.**
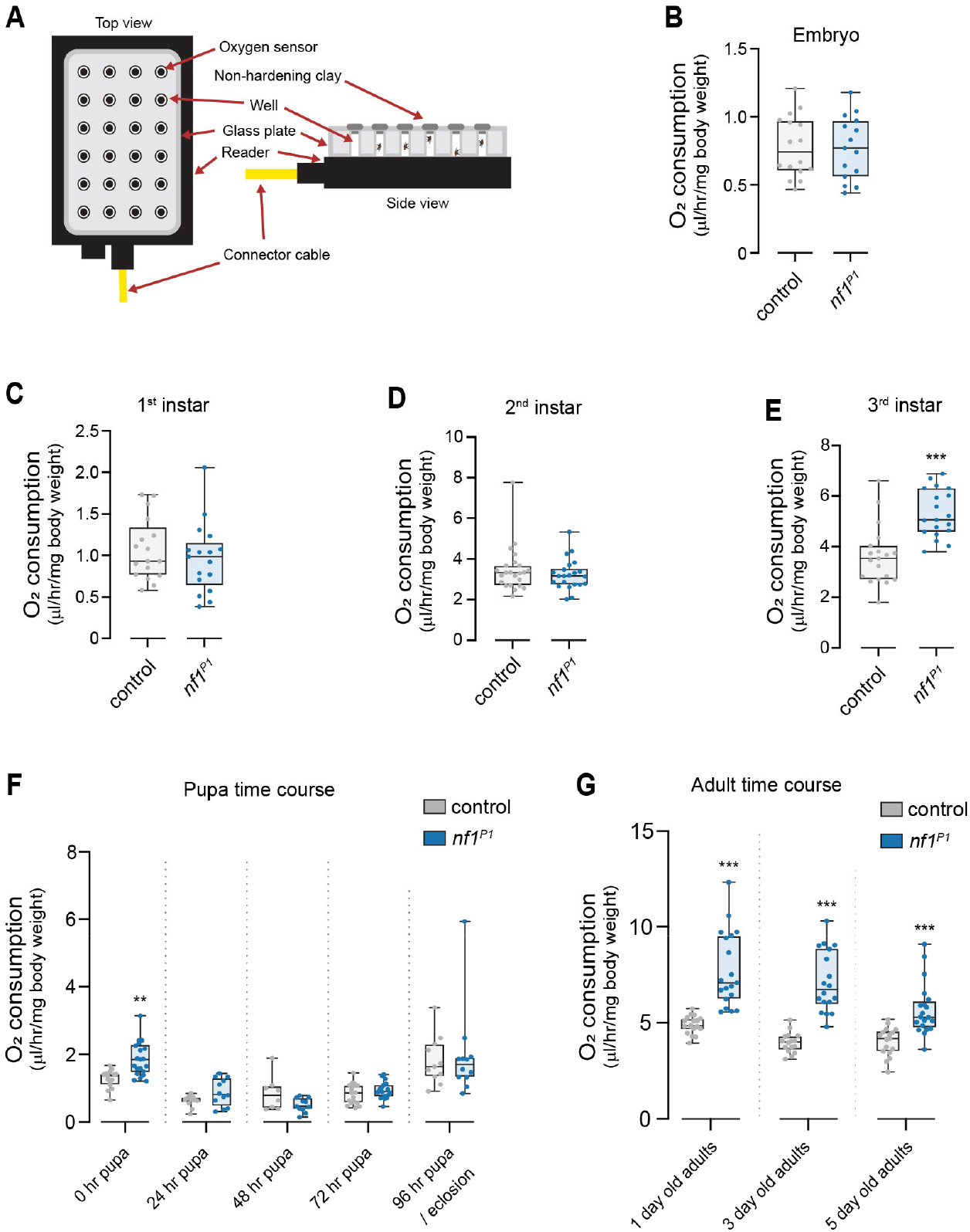
Ontogeny of the Nf1 metabolic effect. **(A)** Diagram of O_2_ consumption respirometry setup. **(B)** O_2_ consumption in *nf1*^*P1*^ mutant embryos, compared to the control line (wCS10). p=0.838 (Mann-Whitney). **(C)** O_2_ consumption in 1^st^ instar *nf1*^*P1*^ mutant larvae, compared to wCS10 controls. p = 0.389 (Mann-Whitney). **(D)** O_2_ consumption in 2^nd^ instar *nf1*^*P1*^ mutant larvae, compared to wCS10 controls. p = 0.521 (Mann-Whitney). **(E)** O_2_ consumption in 3^rd^ instar *nf1*^*P1*^ mutant larvae, compared to the wCS10 controls. ***p < 0.001 (Mann-Whitney). **(F)** Pupal O_2_ consumption time course (0, 24, 48, 72, and 96 hr after pupa formation), comparing *nf1*^*P1*^ mutants to wCS10 controls. **p < 0.01, p = 0.753 for 24 hr, p = 0.611 for 48 hr, p = 0.991 for 72 hr, p = 0.977 for 96 hr (ANOVA/Šidák). **(G)** Adult O_2_ consumption time course (1 day old, 3 day old and 5 day old flies), comparing *nf1*^*P1*^ mutants to wCS10 controls. ***p < 0.001 (ANOVA/Šidák).

To further probe O_2_ consumption, we tested two additional mutants at key time points: 3^rd^ instar larvae, mid- and late-pupal stages (48 and 96 hr), and early adulthood (1, 3, and 5-days post-eclosion). *nf1*^*E1*^ mutant pupae did not exhibit a significant difference in O_2_ consumption rate relative to controls at the 3rd instar larval stage (Fig. 5A) or 48 hr after pupal formation (Fig. 5B). However, by 96 hr after pupal formation, a significant increase in O_2_ consumption was detected (Fig. 5B). After eclosion, there was no significant difference at 1 day post-eclosion, but a significant difference was detected again at 3 and 5 days old (Fig. 5C). Thus, increases in O_2_ consumption began in the pupal phase, and were observed at all time points except the first day post-eclosion. A qualitatively similar pattern was observed with the *nf1*^*C1*^ indel mutant, with some differences at certain time points. This mutant exhibited a significant increase in O_2_ consumption in the 3rd instar larval phase (Fig. 5D); while this differed from *nf1*^*E1*^, it was similar to *nf1*^*P1*^ (Fig. 4E). *nf1*^*C1*^ exhibited no difference in pupal O_2_ consumption at 48 hr after pupal formation, though there was a significant increase relative to controls at 96 hr (Fig. 5E) (similar to *nf1*^*E1*^). After eclosion, *nf1*^*C1*^ adults exhibited no significant difference in O_2_ consumption at 1 and 3 days (though there was a small trend in both) (Fig. 5F). By 5 days post-eclosion, there was a significant increase in O_2_ consumption (Fig. 5F). Since there was some variability across genomic mutations in O_2_ consumption across various time points, we sought to verify the O_2_ consumption phenotype (in adulthood) with a complementary approach, knocking down Nf1 pan-neuronally with RNAi. When Nf1 was knocked down pan-neuronally with RNAi, the experimental knockdown group exhibited a significant increase in O_2_ consumption relative to both controls (Fig. 5G). Integrating the above data, we found no evidence for Nf1 effects on O_2_ consumption prior to the developmental critical period spanning the third instar/pupal phase. The phenotype first emerged in the third instar/pupal phase (O_2_ consumption) or adulthood (CO_2_ production), with the temporal dynamics varying some-what depending on the mutation.

**Figure 5.**
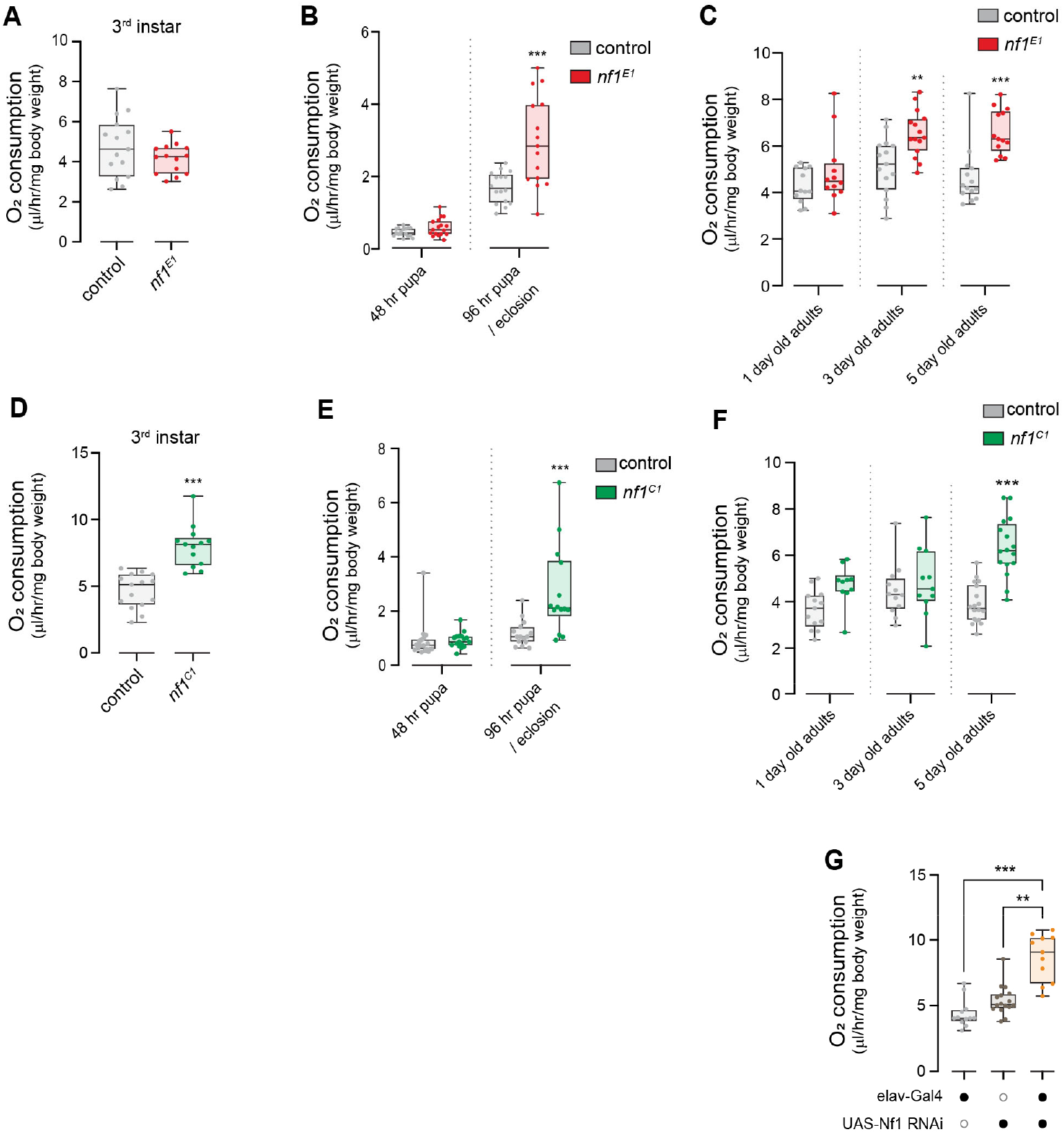
Oxygen consumption in late development and adulthood in *nf1* nonsense mutants. **(A)** O_2_ consumption in 3^rd^ instar *nf1*^*E1*^ mutant larvae, compared to iso2,3 controls. p = 0.377 (Mann-Whitney). **(B)** O_2_ consumption in pupal *nf1*^*E1*^ mutants, compared to iso2,3 controls. p = 0.801 for 48 hr, ***p < 0.001 (ANOVA/Šidák). **(C)** Adult O_2_ consumption time course (1 day old, 3 day old, and 5 day old flies), comparing *nf1*^*E1*^ mutants to iso2,3 controls. p = 0.45 for 1 day old, **p < 0.01, ***p < 0.001 (ANOVA/Šidák). **(D)** O_2_ consumption in 3^rd^ instar *nf1*^*C1*^ mutant larvae, compared to iso2,3 controls. ***p < 0.001 (Mann-Whitney). **(E)** O_2_ consumption in pupal stage *nf1*^*C1*^ mutants, compared to iso2,3 controls. p = 0.996 for 48 hr, ***p < 0.001 (ANOVA/Šidák). **(F)** Adult O_2_ consumption time course (1 day old, 3 day old, and 5 day old flies), comparing *nf1*^*C1*^ mutants to iso2,3 controls. p = 0.069 for 1 day old, p = 0.84 for 3 day old, ***p < 0.001 (ANOVA/Šidák). **(G)** O_2_ consumption in flies with Nf1 knocked down pan-neuronally using elav-Gal4 to drive UAS-Nf1 RNAi, 5 days old. **p < 0.01. ***p < 0.001 (Kruskal-Wallis/Dunn).

### Loss of Nf1 does not detectably alter mitochondrial complex I/II activity or fatty acid oxidation

Alterations in several molecular pathways could lead to alterations in metabolism, including increased O_2_ consumption and CO_2_ production. These include changes in pyruvate oxidation and the tricarboxylic acid (TCA) cycle (in the mitochondrial matrix), and/or the pentose phosphate pathway (PPP) (in the cytoplasm). To test the mitochondrial contributions to changes in metabolism, we isolated mitochondria and carried out high-resolution monitoring of mitochondrial respiratory activity, quantifying leak respiration (LEAK) and oxidative phosphorylation (OXPHOS) flux through electron transport chain complexes I and II. We first compared *nf1*^*P1*^ mutants to K33 controls. In these experiments, K33 were selected as the comparison group to match flies for w+ status [34]. Complex I (CI)-linked respiration was examined by measuring O_2_ flux following the addition of saturating concentrations of malate, glutamate (LEAK), and then ADP (CI-linked OXPHOS). We observed no significant differences between *nf1*^*P1*^ mutants and controls in LEAK or CI-linked respiration (Fig. 6A). Complex II (CII)-linked respiration was examined by measuring O_2_ flux following application of rotenone and succinate. There was a significant increase in O_2_ flux in *nf1*^*P1*^ mutants compared to controls following addition of succinate (Fig. 6A). Following up on this observation in *nf1*^*P1*^ mutants, we tested LEAK and CI/CII-linked respiration in mitochondria isolated from *nf1*^*E1*^ mutants using the same approach. We observed no changes in either CI- or CII-linked respiration (or LEAK) in *nf1*^*E1*^ mutants (Fig. 6B). There are several possible reasons for the difference in observed CII-linked flux between these mutants, including an effect from E(spl)-C genes deleted in *nf1*^*P1*^ mutants and/or an effect from gut microbiota potentially co-isolated with mitochondria. To test for potential nonspecific effects of gut microbiota, we tested mitochondria samples isolated from a cohort of *nf1*^*P1*^ mutants and controls following streptomycin feeding. There was no difference in LEAK, CI- or CII-linked respiration in these samples (Fig. 6C). Next we examined fatty acid oxidation by supplying substrates for β-oxidation, palmitoylcarnitine, malate, and ADP, in *nf1*^*E1*^ mutants. There was no difference in O_2_ flux between *nf1*^*E1*^ mutants and controls (Fig. 6D). Overall, these data suggest that there was neither a detectable change in mitochondrial respiration through complex I or II pathways nor through fatty acid oxidation in adult *nf1* mutants. Thus, the adult metabolic alteration in Nf1 loss of function conditions likely occurs upstream of the mitochondrial electron transport chain.

**Figure 6.**
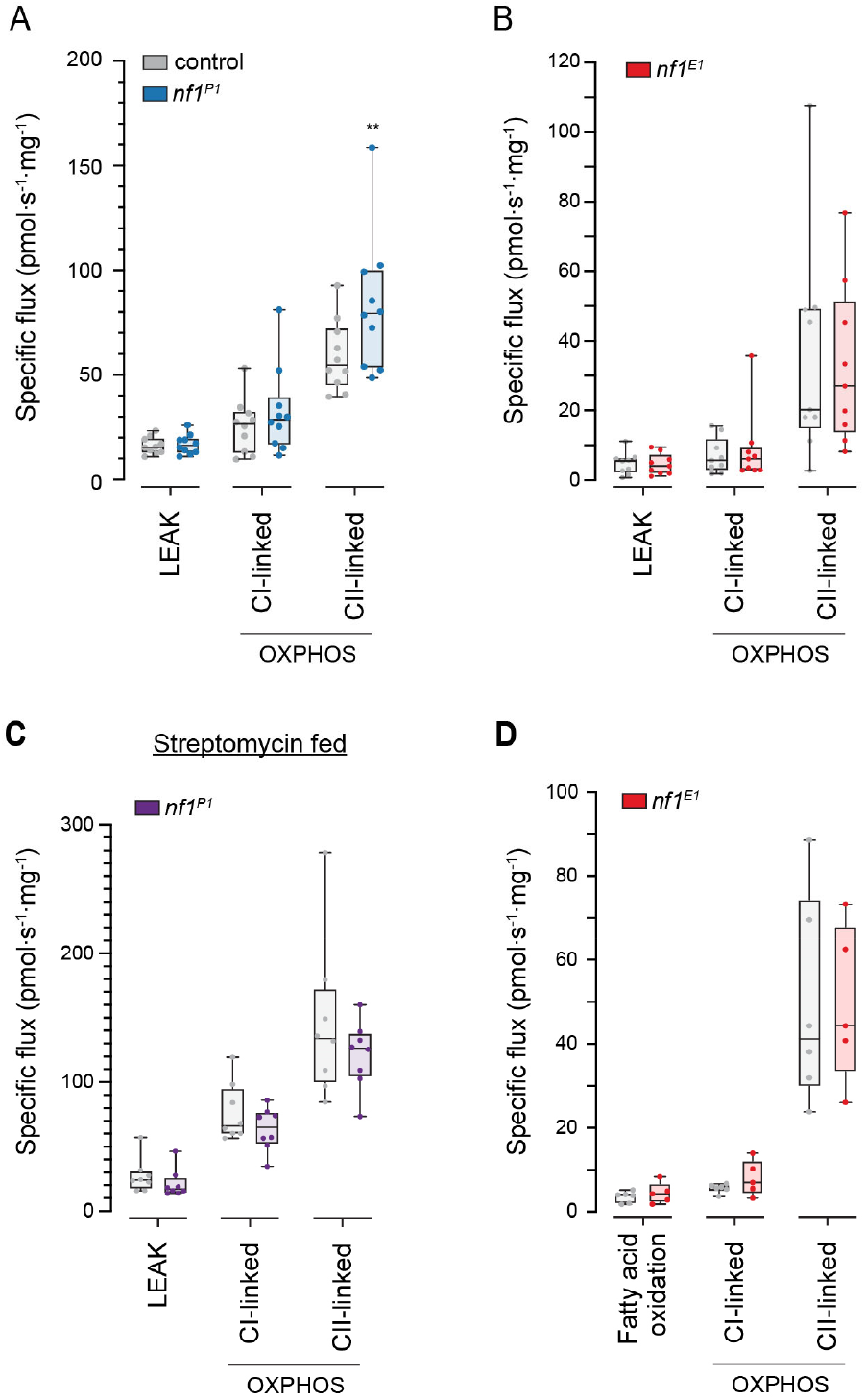
Nf1 deficiency does not detectably alter mitochondrial CI/CII-linked respiration or fatty acid oxidation. **(A)** High-resolution respirometry comparing LEAK, complex I (CI) and complex II (CII)-linked respiratory activity between mitochondria isolated from *nf1*^*P1*^ mutants and those from K33 controls. **p < 0.01 (ANOVA/Šidák). **(B)** Respirometry in mitochondria isolated from *nf1*^*E1*^ mutants compared to those from iso2,3 controls. **(C)** Respirometry in mitochondria isolated from *nf1*^*P1*^ mutants compared to those from K33 controls, both fed streptomycin. **(D)** Fatty acid oxidation (FAO) + CI + CII-linked respiration in *nf1*^*E1*^ mutants and controls.

## Discussion

Neurofibromatosis type 1 drives a range of symptoms, including some that have neurodevelopmental contributions, such as attention-deficit/hyperactivity disorder and autism spectrum disorder [1, 20, 35, 36]. Genetic mutations in the NF1 gene alter the activity of signaling pathways such as Ras/MAPK, which influence cellular growth and differentiation. In addition, the disease produces metabolic shifts, which include changes in resting energy expenditure [10], lower fasting blood glucose and increased insulin sensitivity [9], changes in body composition [8] including reduced triglyceride levels [37], and reduced incidence of diabetes mellitus [1, 38–40]. The mechanisms underlying these metabolic alterations are not known, and it is unclear whether they are interrelated with the neurodevelopmental symptoms.

Animal models have suggested an Nf1-dependent developmental contribution to adult phenotypes. Loss of Nf1 increases spontaneous grooming frequency in *Drosophila*, a repetitive behavior that is reminiscent at the behavioral level of hyperactivity/ADHD symptoms in humans [24, 41]. The effect is robust, with *nf1* mutants spending up to 7x more time grooming than control flies. It is developmental in origin – loss of Nf1 during a developmental critical period produces abnormal grooming patterns, while loss of Nf1 in adulthood does not [21]. In a conceptually similar manner, loss of Nf1 alters signaling pathways in a way that may influence cancer susceptibility via developmental mechanisms. For instance, MEK inhibition treatment during a developmental window prevents optic glioma formation via ERK-dependent effects on migrating glial progenitors [42].

Here we found that the Nf1 metabolic phenotype is also subject to developmental control. Loss of Nf1 during a developmental critical period drives the adult phenotype, and the critical period for metabolic dysregulation matches the critical period for behavioral regulation (grooming) [21]. Both the metabolic phenotype and grooming phenotype result from loss of Nf1 in neurons [12, 21, 24]. The neuronal circuits that regulate the grooming and metabolic phenotypes are distinct [12, 43], suggesting that common neurodevelopmental processes can drive multiple organismal endpoints by affecting different neuronal circuits. Such processes may include alterations in fundamental neuronal functions such as developmental regulation of neuronal excitability [44], synaptic development/refinement [45], synaptic transmission/plasticity, neuromodulatory circuit function [46, 47], or inhibitory circuit function [48–50]. Each of these could affect multiple circuits and downstream organismal behavioral phenotypes. Dissecting the neurodevelopmental mechanisms that drive adult phenotypes will be a key area of investigation for further mechanistic analysis.

The developmental contribution of Nf1 to adult metabolism was unexpected, despite its developmental roles in other contexts. Nf1 impacts signaling cascades that directly affect mitochondrial function in adult animals, suggesting dynamic real-time metabolic modulation. For instance, in an oncogenic context, Nf1 regulates mitochondrial function via ERK-mediated phosphorylation of the chaperone protein TRAP1 [51]. Whether this metabolic function of Nf1 operates in parallel in a non-oncogenic context is not known. Yet we did not detect differences in mitochondrial complex II function. This suggests that there may be a divergence in the primary mechanism of Nf1 neurodevelopmental function, as TRAP1 phosphorylates succinate dehydrogenase (complex II).

Prior studies established that loss of Nf1 alters metabolic function in adult *Drosophila* [1, 12] and suggest that mitochondrial function could be altered. Metabolic alterations include increased ROS production and reduced lifespan [13], increased O_2_ consumption and CO_2_ production with reduced respiratory quotient [12, 21, 24, 50], reduced triglyceride stores and increased rate of lipid turnover [12], reduced glycogen stores [52], increased feeding [12], reduced growth and body size [14, 26, 53], sleep-metabolism interaction [50], elevated NADase SARM1 levels [25], and changes in mitochondrial morphology [25, 43]. We observed no significant differences in mitochondrial electron transport chain complex I/II function in mitochondria isolated from *nf1* mutants, suggesting that the metabolic alterations are independent of - and likely upstream of - the electron transport chain. Given that neuronal loss of Nf1 drives the phenotype, neuronal Nf1 could impact metabolism in either cell-autonomously (in neurons) and/or non-cell-autonomously (in peripheral tissues) via neurodevelopmental effects.

Several pharmacological treatment strategies have been developed for cognitive and behavioral symptoms of neurofibromatosis type 1 have been tested in humans, including antiseizure drug lamotrigine and the HMG-CoA reductase inhibitors lovastatin and simvastatin. These treatments have shown limited efficacy in alleviating cognitive and behavioral symptoms in clinical trials [16–20]. The relative lack of effect may lie with selection of targets, compounds, or assessment of cognitive and behavioral measures. Prior studies and the present study suggest that neurodevelopmental effects of Nf1 contribute to development of multiple phenotypes in adult animals [21]. Basic neurodevelopmental processes are conserved across taxa, and, in addition to Nf1 itself, mutations in signaling pathways downstream of Nf1 drive neurodevelopmental alterations [22]. If these findings translate to humans, then the timing of drug intervention may prove critical for treating cognitive and behavioral symptoms in neurofibromatosis type 1 and intervening earlier in life is likely to be necessary.

Neurodevelopment in humans may be affected in neurofibromatosis type 1, which alters corpus callosum morphology and optic nerve tortuosity [1]. In addition, there are changes in neuronal activity, including functional connectivity within the default network, corticostriatal circuits, and other regions [1]. Some of these changes are recapitulated in animal models of neurofibromatosis type 1. For instance, conditional knockout of Nf1 in mice enlarges the corpus callosum, similar to humans with the disorder [54]. This is developmental, as inhibiting ERK during neonatal development restores normal corpus callosum volume [54]. Thus, Nf1 deficiency changes brain structure and function in humans, and animal models have implicated neurodevelopmental processes in some of these.

Which molecular signaling pathways downstream of Nf1 (e.g., MAPK, mTOR, Ral, etc.) are primarily responsible for the developmental effects of Nf1 are not known. Current treatments for NF1 (cancers) focus on MEK, and several studies implicate downstream ERK in developmental phenotypes [22, 54]. Yet other pathways may also be involved. Nf1 is a Ras GAP, which reduces Ras activity, in turn affecting multiple signaling pathways downstream of Ras [1]. These include direct down-stream effectors such as MAPK, mTOR, Ral, and other pathways that are indirectly modulated, such as cAMP/PKA. There is significant crosstalk between these signaling pathways [43, 55, 56]. In addition, there are multiple biochemical routes through which they could affect neuronal function and metabolic processes [51, 57–59]. In *Drosophila*, both the grooming [21] and metabolic phenotypes (present study) are dependent on Nf1 effects in neurons during development [1, 12, 24], though they map to dissociable neuronal circuits [12]. Which signaling pathways downstream of Nf1 are responsible and their cellular targets/mechanisms are not fully elucidated.

Whole-animal CO_2_ production was increased in adult *Drosophila* with Nf1 deficiency, a finding reported previously [21, 24, 50], and identified here as a developmental phenomenon. There are three major metabolic sources of CO_2_: pyruvate oxidation, the TCA cycle, and the PPP. The first two are carried out in the mitochondrial matrix, while the latter occurs in the cell cytosol. In search of the mechanism of the increased CO_2_ production, we carried out high-resolution respirometry of mitochondria, finding no detectable alteration of mitochondrial complex I/II function. Thus, the increase in CO_2_ production may occur upstream of the electron transport chain, and/or involve the PPP pathway. The increase in O_2_ consumption points primarily to changes in mitochondrial number or electron transport chain function. In addition, peroxisomal respiration contributes up to 20% of total O_2_ consumption [60], and could therefore be involved. Further studies will be necessary to determine the cellular mechanism for the increase in O_2_ consumption and CO_2_ production.

Results from the present study raise a “chicken or the egg” causality question in terms of neurodevelopmental and metabolic effects in Nf1 deficiency. The metabolic phenotype is due to loss of Nf1 in neurons [12]. In this study, we found that it is due to loss of Nf1 in neurons specifically during a developmental critical period. This is the same critical period for neurodevelopmental effects on at least one behavioral phenotype [21]. One possibility is that loss of Nf1 alters neurodevelopment, which subsequently rewires systemic metabolic control via central circuits. Alternatively, loss of Nf1 could alter metabolism in a cell-autonomous fashion first, changing developmental trajectory via effects on cellular proliferation, differentiation, migration, and/or maturation processes [61–63]. The signaling pathways altered by Nf1 deficiency, particularly Ras/ERK, could potentially influence either/both of these processes (neurodevelopment and stem cell metabolism) in various ways.

Overall, this study revealed that metabolic changes with loss of Nf1 in *Drosophila* are due in large part to neurodevelopmental effects. Loss of Nf1 during a developmental critical period alters metabolism in adult animals. This is reflected in increased O_2_ consumption and CO_2_ production in the whole animal, which emerge during/after the critical period. These findings suggest that the metabolic shifts in neurofibromatosis type 1, like the cognitive and behavioral phenotypes, may exhibit dependence on developmental processes.

## Acknowledgements

We thank Aaron Stahl for feedback on the experiments and manuscript draft. We thank Andre Bernards for *nf1*^*P1*^ fly stocks, James Walker for *nf1*^*E1*^, *nf1*^*C1*^, UAS-Nf1, and iso2,3 fly stocks, and Ronald L. Davis for wCS10 fly stocks. Additional stocks used in this study were obtained from the Bloomington *Drosophila* Stock Center (NIH P40OD018537) and the Vienna *Drosophila* Resource Center. We thank Eric Weatherford and the University of Iowa Metabolic Phenotyping Core Facility for assistance with Oroboros O2k analysis. This research was supported by the National Institutes of Health R01 NS114403 to SMT, R01 NS124716 to SMT, R01 NS126361 to SMT, R01 NS097237 to SMT, Department of Defense NF230039 to SMT, and an award from the Roy J. Carver Charitable Trust to SMT. We thank Linda Buckner and Robert Svetly for administrative assistance. This preprint was typeset with the bioRxiv word template by @Chrelli: www.github.com/chrelli/bioRxiv-word-template

## Materials and Methods

### Fly maintenance and genetics

Flies were raised on cornmeal/agar food medium according to standard protocols. Flies were housed in incubators at 25°C, 60% relative humidity and on a 12:12 hr light:dark cycle. Genomic mutants were compared to controls with matched genetic backgrounds (wCS10 and K33 for *nf1*^*P1*^, iso2,3 for *nf1*^*E1*^ and *nf1*^*C1*^). The Nf1 RNAi line was obtained from the Vienna Drosophila Resource Center (VDRC #109637) [28]. UAS-dicer2 was used in all crosses to enhance the RNAi effect [28]. Gal4/+ control crosses contained an empty attP control line (VDRC #60100) to ensure a matched genetic background across all groups. Male flies were used for all experiments, unless otherwise specified. Flies were collected at specific ages, including embryo, larvae (1st, 2nd, and 3rd instar), pupae (0, 24, 48, 72, and 96 hr after pupal formation), and adult (1, 3, and 5 days after eclosion). Larval stages were ascertained by counting cephalopharyngeal sclerites. Non-staged adult flies (Fig. 1) were collected from 5-7 days old. Staged adult flies were collected over an 8 hr window (1 or 3 days old) or 24 hr window (5 days old).

### Streptomycin feeding

Streptomycin was dissolved in water and added to the cornmeal/agar fly food to a final concentration of 400 μg/mL [64–66]. Flies were fed on standard food from embryo through pupariation and transferred to streptomycin-containing food following eclosion.

### Conditional Nf1 knockdown

UAS-Nf1 RNAi was driven pan-neuronally by elav-Gal4 under temperature-sensitive control of tub-Gal80ts. The experimental genotype was: elav-Gal4/+;tub-Gal80ts/UAS-Nf1RNAi;UAS-dcr2/+. Non-RNAi expressing temperature controls were run in parallel consisting of the driver/repressor driving UAS-GFP: elav-Gal4/+;tub-Gal80ts/UAS-GFP. Nf1 was knocked down during early development by raising flies at 30 °C from embryo through 2^nd^ instar larvae, then switching them to 18 °C from the 3^rd^ instar larvae through adult stage. For late development knockdown, flies were maintained at 18 °C from embryo through the 2^nd^ instar stage, shifted to 30°C during the 3^rd^ instar and pupal stages, and returned to 18 °C at eclosion. For adult-specific knock-down, flies were raised at 18 °C from embryo through the pupal stage, then transferred to 30 °C following eclosion. Negative control flies were kept at 18 °C from embryo through adulthood. The genetic background line and heterozygous controls were tested with the late development temperature shift (and no shift).

### Carbon dioxide respirometry

CO_2_ production was measured with respirometry [67]. Respirometers were created by gluing a 1 mL pipette tip and a 50 µL capillary micropipette together. Sixteen respirometers were hung on a custom-made rack in a latch-lid chamber filled with a red water-based dye solution. Soda lime was placed into each pipette tip between two foam pieces. CO_2_ anesthesia was used to sort flies and allowed 24 hr for flies to recover before beginning the experiment. Four flies of the same genotype were placed into each pipette using an aspirator and one respirometer was left empty during each experiment as a temperature and atmospheric control. The top of the pipettes were tightly sealed using non-hardening modeling clay. Respirometers that were not tightly sealed were excluded from final analysis. The chamber with the flies were left to equilibrate to incubator conditions (25°C) for 1 hr before starting the experiments, and vacuum grease was used to seal the chamber lid to reduce atmospheric and temperature fluctuations. Images were captured every 15 min with PhenoCapture 3.3. The liquid meniscus level in each respirometer after 3 hr was measured using Fiji 2.0.

### Oxygen microplate respirometry

Whole-animal O_2_ consumption was measured using a 24-well microplate respirometer with optical O_2_ sensors (Loligo Systems, Denmark) [67]. 200µL well plates were used for adult *Drosophila* and 80µL well plates were used for embryos, larvae, and pupae. All experiments were performed at room temperature (21.5 ± 2 °C), and microplates were calibrated using two-point calibration, room air for high calibration (100% air saturation) and 100% N2 for low calibration (0% air saturation). Wells were closed with non-hardening modeling clay, with 4 wells per plate left empty to record background changes in O_2_. The first 30 min of each run were excluded from analysis as an incubation duration to allow animals to adjust to the experimental set up. O_2_ consumption was calculated based on the linear decline in O_2_ concentration in the wells over a 3 hr time period and subtracting the blanks from the samples’ values. Values with r2 < 0.9 were excluded from analysis.

### High-resolution mitochondrial oxygen respirometry

Mitochondrial preparations were obtained from 5-6 day old *Drosophila melanogaster* by homogenization in ice-cold MiR05 respiration buffer (0.5mM EGTA, 3mM MgCl2, 60mM Lactobionic acid, 20mM Taurine, 10mM KH2PO4, 20mM HEPES, 110mM D-Sucrose, 1g/L fatty acid free bovine serum albumin, pH 7.1) and two rounds of centrifugation at 700xg at 4°C for 5 min to remove cellular debris. Male and female flies were combined in each replicate. Isolated mitochondria were then added to 2mL chambers of an O2k high-resolution respirometer (Oroboros Instruments, Innsbruck, Austria). Experiments were run at 30°C in MiR05 respiration buffer. Mitochondrial respiration was quantified by measuring O_2_ consumption rates following a sequential substrate inhibitor titration protocol using DatLab8 (Oroboros Instruments). Titration of malate (2 mM) and glutamate (10 mM) stimulated non-phosphorylating respiration through complex I. Oxidative phosphorylation capacity was measured following titration of 1.25 mM ADP. Next, complex I respiration was inhibition by rotenone (0.5 μM) and succinate was added to stimulate complex II respiration. Antimycin A (2.5 μM), a complex III inhibitor, was added to evaluate residual O_2_ consumption and as the baseline flux [69, 70]. In separate experiments, fatty acid oxidation (FAO)-dependent mitochondrial respiration was measured using 0.1mM malate, 10uM Palmitoyl-L-Carnitine and 1.25mM ADP. Malate (2 mM) and glutamate (10 mM) were added to stimulate the complex I (CI) respiration (FAO & CI pathway). Rotenone (0.5 μM) and Succinate (10 mM) were added to measure complex II capacity. Antimycin A (2.5 μM) was used to evaluate the residual O_2_ consumption and as the baseline flux [71]. Respiratory flux was normalized to protein concentration. Protein concentration was calculated using Pierce BCA Protein assay [69, 70].

### Statistical analysis

Normality of data was assessed with the D’Agostino Pearson Test. In figures, box plots graph the median as a line, the interquartile range (IQR) as a box, and whiskers extend to the min/max values. Hypothesis testing was carried out using t tests or ANOVA followed by Dunnett (comparison with one group) or Šidák’s (comparison with multiple groups) multiple comparisons tests (parametric), or the Mann-Whitney U test or Kruskal-Wallis omnibus test followed by Dunn multiple comparisons tests (nonparametric). For RNAi analysis, the experimental group was compared to heterozygous Gal4/+ and UAS/+ controls and considered positive only if it significantly differed from both controls in the same direction. Statistics and graphing were carried out with Graphpad Prism, version 11.0.0.

